# Filamentation profile reveals several transcription regulators that contribute to differences between *Candida albicans* and *Candida dubliniensis*

**DOI:** 10.1101/2024.11.20.624441

**Authors:** Teresa Meza-Davalos, Luis F. García-Ortega, Eugenio Mancera

**Author notes:** Correspondence to Eugenio Mancera.

## Abstract

*Candida dubliniensis* is the most closely related species to *C. albicans,* one of the leading causes of fungal infections in humans. However, despite sharing many characteristics, *C. dubliniensis* is significantly less pathogenic. To better understand the molecular underpinnings of these dissimilarities, we focused on the regulation of filamentation, a developmental trait fundamental for host colonization. We generated a collection of 44 *C. dubliniensis* null mutants of transcription regulators whose orthologs in *C. albicans* had been previously implicated in filamentous growth. These regulators are very similar at the sequence level, but phenotypic screening identified several mutants with contrasting interspecific filamentation phenotypes, beyond previously known differences. Bcr1, a well-known regulator of biofilm formation, stands out as its mutant only showed a filamentation defect in *C. dubliniensis*. Phenotypic and transcriptional characterization showed that the *bcr1* defect is condition dependent and that this regulator plays a central role in the filamentation of *C. dubliniensis*, possibly by regulating the hyphal activator Ume6. Overall, our results suggest that several regulatory pathways are involved in the filamentation differences between *C. albicans* and *C. dubliniensis* and show that the *C. dubliniensis* mutant collection is a valuable resource to compare, at a molecular level, these two species of medical relevance.

**AUTHOR SUMMARY:** The yeast *Candida albicans* is one of the most important fungal pathogens for humans. Its ability to form filamentous cells is central for the colonization of the human body. Although *Candida dubliniensis*, the closest known relative to *C. albicans*, is also able to filament, it is a much rarer cause of disease. Part of the virulence differences between these species have been attributed to their filamentation dissimilarities, but we are just starting to understand the regulatory pathways that control filamentation in *C. dubliniensis*. Here, we generated a collection of gene-deletion mutants in *C. dubliniensis* of the orthologs of transcription regulators that have been associated with filamentation in *C. albicans*. Comparative profile of the collection revealed that several regulators contribute to the filamentation dissimilarities between the two species. Among these, our results identified Bcr1 as a regulator with a prominent role controlling filamentation in *C. dubliniensis*, showing that its target genes have considerably changed between *C. albicans* and *C. dubliniensis*. Our findings and the collection of mutants that we generated open new opportunities to better understand the molecular mechanisms that underlie the pathogenicity of these clinically important microorganisms.

## INTRODUCTION

Fungi from the genus *Candida* are among the most important human pathogens (Bennett, 2010). They are capable of causing a spectrum of diseases, ranging from mild superficial infections of the oral cavity and vagina to severe, life-threatening systemic conditions with high levels of morbidity and mortality (1, 2). These infections pose a significant risk especially to individuals with compromised immune systems (1). However, these fungi are also found as part of the microbial communities that commensally inhabit our bodies and therefore are considered opportunistic pathogens.

Most medically relevant *Candida* species belong to the CTG (CUG-Ser1) clade, a monophyletic group of ascomycetous yeasts characterized by the translation of the CUG codon as serine instead of leucine (3). Within this group, *Candida albicans* stands out as the most virulent species, being a leading cause of both superficial and systemic infections in humans (4). The clade includes other important opportunistic pathogens, but also many species that have not been associated to humans or that are much rarer ethological agents. This is the case with *C. dubliniensis*, the species most closely related to *C. albicans* phylogenetically, yet considerably less prevalent in clinical settings. For example, *C. albicans* has been estimated to be responsible for approximately 65% of the infections caused by *Candida* species, while *C. dubliniensis* accounted for fewer than 0.1% (5). In agreement, *C. dubliniensis* has been shown to be less virulent in several murine models of infection (6-8). Given their virulence differences but evolutionary proximity – the two species are estimated to have diverged 20 million years ago (9, 10) – *C. dubliniensis* has been a useful comparative model to understand the underpinnings of *C. albicans* pathogenicity (2).

In *C. albicans,* the morphological transition between yeast and filamentous cells (hyphae and pseudohyphae) is important for the colonization of the human host and disease causation (11). Changes in cell-shape have been suggested to allow disruption of host cells and tissue, while the differential expression of virulence factors between yeasts and filaments is also key for the adaptation to different environments in the host, including interactions with other microorganisms in these habitats (12). The ability to filament is shared by *C. dubliniensis*, as this species is also able to form both hyphae and pseudohyphae. However, relative to *C. albicans, C. dubliniensis* has been observed to filament more rarely and the range of known stimuli that trigger filament formation in this species is much narrower. Not surprisingly, the decreased ability to transition from yeast to hyphae has been associated with its reduced virulence (6-8). In agreement, experiments in murine models of infection have shown that *C. dubliniensis* cells in the stomach and kidney remain predominantly in the yeast form, whereas *C. albicans* cells exhibited both yeasts and hyphae (7, 8).

Multiple signaling pathways and transcriptional regulators have been found to control the switch between yeast and filamentous forms in *C. albicans,* indicating that the morphological transition is quite complex at a molecular level (13, 14). Changes in some of these pathways and regulators have been associated with the filamentation differences with *C. dubliniensis*. For example, differential expression of the transcriptional repressor Nrg1 has been shown to be partially responsible for the filamentation differences as it is quickly downregulated in *C. albicans* by several stimuli of the human host, while its expression does not decrease as sharply in *C. dubliniensis* (15). Similarly, overexpression of the transcription regulator Ume6 that is known to be repressed by Nrg1 (15, 16) has been associated with filamentation of *C. albicans* under several conditions, but in *C. dubliniensis* its expression change requires starvation, one of the few conditions where this species is known to filament (17). Although the genomes of *C. albicans* and *C. dubliniensis* are very similar, their comparison also shed light into the filamentation differences of the two species (2). For instance, key hypha-specific virulence factors such as Hyr1, Als3 and some members of the secreted aspartyl proteinase (SAP) family are absent in *C. dubliniensis* (17, 18). These differences have also been observed when comparing genome-wide transcriptional profiles of the two species, revealing hypha induced genes in *C. albicans* that do not change their expression in *C. dubliniensis* (18).

Given the complexity of the regulatory circuit that controls filamentation in *C. albicans* – at least 45 transcription regulators have been associated with this transition – additional differences with *C. dubliniensis* could be expected. To further understand the dissimilarities at a molecular level, we generated a deletion collection of most of the *C. dubliniensis* orthologs of the transcription regulators that have been associated with filamentation in *C. albicans.* Comparative profiling under inducing conditions revealed contrasting filamentation phenotypes in several of the mutants, beyond the previously known differences. Transcriptional profiling of the *bcr1* mutant, one of the regulators with marked differences, showed extensive interchange of target genes. Overall, our work suggests considerable rewiring in the regulatory circuits that control filamentation in these two closely related species with contrasting clinical characteristics.

## RESULTS

### TRs that control filamentation are conserved at the sequence level between *C. albicans* and C. dubliniensis

To better understand the differences in the molecular mechanisms that control filamentation between *C. albicans* and *C. dubliniensis* we focused on the TRs that have been associated with this cellular process. TRs function as hubs in the regulation of cellular metabolism and given the gene deletion collection available for their study in *C. albicans*, they represented an ideal entry point. We found 45 TRs whose knockout mutant in *C. albicans* had a phenotype associated with filamentation in CGD (Materials and Methods; Supplementary Table 2). These TRs represent between 15 and 20% of the total of TRs present in *C. albicans*, and all have a one-to-one ortholog in *C. dubliniensis* according to CGOB (19). To assess the degree of conservation of the 45 TRs between *C. albicans* and *C. dubliniensis*, we first aligned the protein sequence of each pair of orthologs. The average sequence identity between orthologs is of 81.1% and, as expected, it is even higher in the DNA binding domain (96.3%). At the amino acid sequence level, this group of TRs is not atypical since their identity falls well within the range of identity of all the TR ortholog pairs between these two species (Figure 1). This is also the case for the DNA binding domain. In both species, the protein domains that are most common among the 45 filamentation TRs are the Zinc finger C2H2-type (IPR013087), the Zn(2)Cys(6) fungal-type DNA-binding domain (IPR001138) and the Myc-type, basic helix-loop-helix (bHLH) domain (IPR011598), while in the overall set of TRs the most frequent ones are the Zn(2)Cys(6) fungal-type DNA-binding domain (IPR001138), the Zinc finger C2H2-type (IPR013087) and the Transcription factor domain, fungi (IPR007219). The conservation in protein domain composition in the TRs of *C. albicans* and *C. dubliniensis* is in agreement with the high sequence identity of the TRs between the two species.

**Figure 1.**
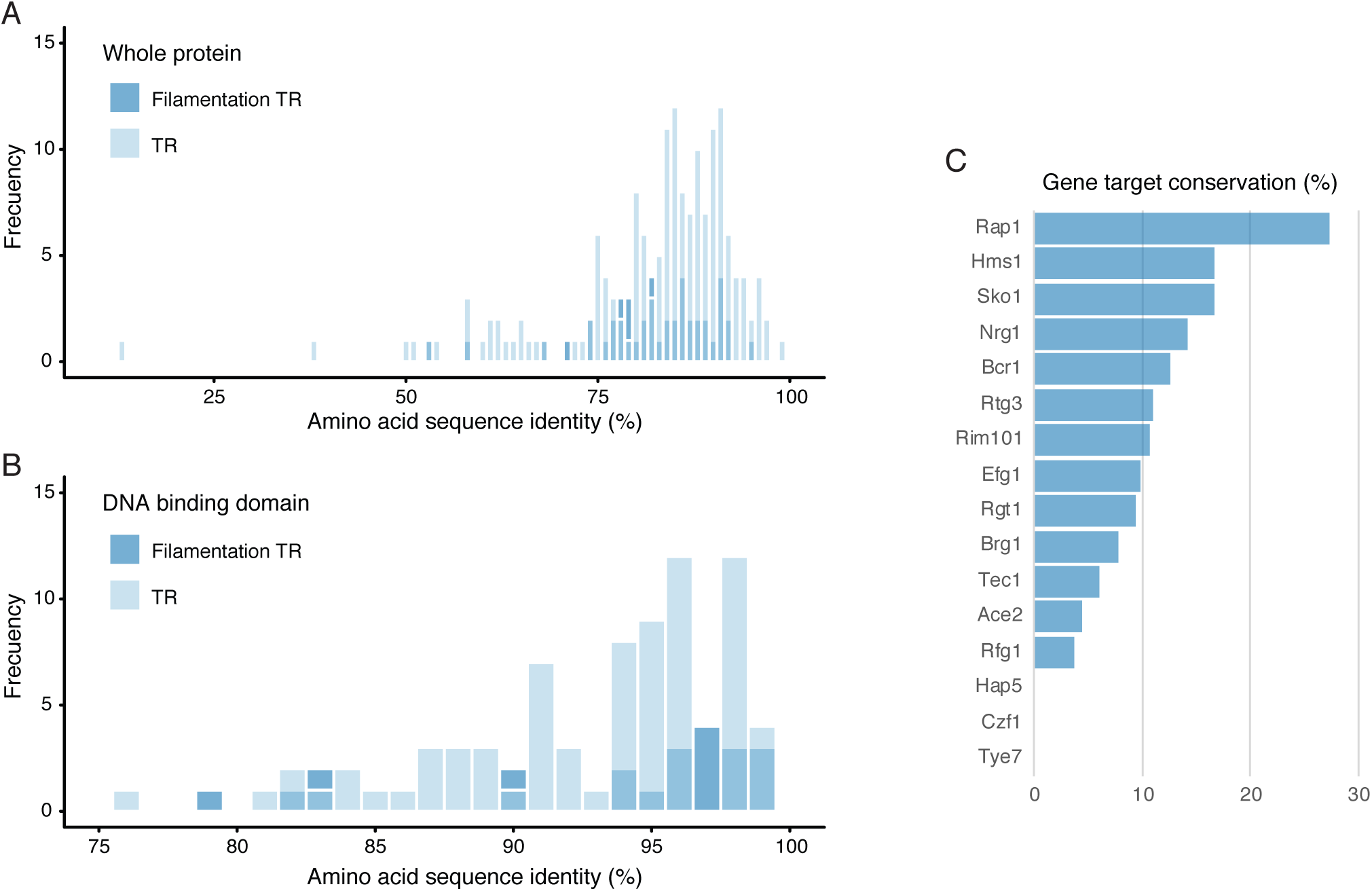
Filamentation TR are conserved at the protein level between *C. albicans* and *C. dubliniensis*, but their putative target genes have diverged considerably. (A) Distribution of the amino acid sequence identity (%) of all the TR orthologs (light blue) and filamentation TR orthologs (dark blue) along the whole protein. (B) Distribution of the amino acid sequence identity (%) of all the TR orthologs (light blue) and filamentation TR orthologs (dark blue) in the DNA binding domain. Only the TR for which a DNA binding domain has been defined are included. (C) Conservation (%) of computationally predicted target genes between filamentation TR orthologs. Target genes were defined by the presence of the DNA binding sequence motif in the upstream region of a gene. Only TR for which a DNA binding sequence motif has been previously reported in *C. albicans* were included.

### The predicted target genes of the filamentation TRs are very different between *C. albicans* and *C. dubliniensis*

Contrary to the high conservation that we observed at the amino acid sequence level, previous experimental determination of the target genes of six of these TRs has shown considerable divergence between *C. albicans* and *C. dubliniensis* (20). To further explore the degree of conservation in target genes, we computationally identified targets based on the presence of the DNA binding motif in their upstream region. The motifs of only 16 TRs have been determined in *C. albicans* and we assumed that these motifs are conserved in *C. dubliniensis* given the sequence similarity between the ortholog TRs, especially in the DNA binding domain. The degree of conservation that we observed in putative target genes fell within the range of previous comparisons between these two species (20). The TRs whose putative target genes are most conserved is Rap1, and only 27% of its targets are shared between *C. albicans* and *C. dubliniensis* (Figure 1). On the other end of the distribution, for 3 TRs (Czf1, Hap5 and Tye7) there were no putative gene targets shared between the two species. Overall, our observations suggest that the filamentation differences between *C. albicans* and *C. dubliniensis* could be due to the target gene differences of the TRs that regulate the process. In addition, they showed that without further experimentation it would be difficult to predict the specific TRs that contribute to the filamentation differences between the two species.

### A TR mutant collection to identify differences in the regulation of filamentation between *C. albicans* and *C. dubliniensis*

To experimentally identify differences between *C. albicans* and *C. dubliniensis* in the function of the TRs that are involved in filamentation, we generated a gene knockout collection of the orthologs of the *C. albicans* filamentation TR in *C. dubliniensis*. Of the 45 filamentation TRs in *C. albicans*, a knockout mutant had already been generated in *C. dubliniensis* for five of them (20). A *nrg1* null mutant had also been generated, but in a different genetic background (15) and therefore we also constructed the deletion strain. The null mutants for the 40 TRs were generated using the same genetic engineering strategy used to generate the *C. albicans* TR mutants, employing two auxotrophic markers to tandemly delete the two alleles of a given gene (21). For each gene, one independent homozygous mutant was generated in two different *C. dubliniensis* parental auxotrophic strains so that the phenotype of the deletion could be assessed in replicates.

Even though several transformation attempts were performed we could not generate the homozygous *tup1* mutant. This was surprising given that deletion mutants of this gene have been previously reported in *C. albicans* and *C. tropicalis* (22, 23). It is possible that this TRs is essential in *C. dubliniensis* or that its gene is located in an aneuploid genomic region, and an extra allele is present in this species. Together with the 5 deletion mutants previously generated, we put together a collection of 44 homozygous and 44 heterozygous knockout mutants to experimentally assess the function of the *C. dubliniensis* orthologs of the filamentation TRs.

### Multiple TRs contribute to the filamentation differences between *C. albicans* and *C. dubliniensis*

To test wheatear the deletion mutants of the TRs have a filamentation defect we performed filamentation time courses of all the mutants in both species in parallel. As detailed in Materials and Methods, filamentation was induced transferring strains to water with 10% fetal bovine serum at 37 °C since both species are known to from hypha under these conditions. Morphological changes were monitored under the microscope at the time of transferring to the induction media (“zero” time-point) and after one and three hours of induction. Filamentation was quantitatively estimated as the fraction of cells that formed a germ tube or hyphal morphologies at each time point. To clarify phenotypic inconsistences between the two isolates of the mutants of five genes (*PHO4, GRF10, AFT2, ADR1* and *FGR15*), we generated an additional homozygous mutant.

We could not quantitate filamentation in all time points for 12 TR mutants (*ace2, czf1, efg1, fgr15, nrg1, rap1, rca1, rfg1, rfx2, ssn6, stp2 and tye7*) given that they showed morphologies that could not be clearly classified as yeast cells or germ tubes in the two species (Supplementary Figure 1). For all of these 12 mutants but *rfg1* the aberrant morphology was already evident in one of the two species at the time of transferring the cells to the induction media, suggesting that the morphology is independent of the filamentation stimulus. Most of these mutants formed elongated cells or pseudohyphal aggregates not observed in the wildtype strains at time point zero. The mutants *ace2, fgr15, rap1, rca1, ssn6* and *stp2* showed deviant morphology in both species and that are in agreement with previous reports in *C. albicans* (4, 15, 21, 24-26). On the other hand, *tye7*, *czf1*, and *rfx2* exhibited pseudohyphal growth in *C. dubliniensis*, while maintaining a yeast form in *C. albicans*. Additionally, the mutant *nrg1* displayed hyperfilamentous growth solely in *C. albicans*. The *efg1* mutant did not filament in any of the three time points considered, however it formed elongated cells, particularly in *C. dubliniensis*. Even when these cells were clearly not germ tubes, they made quantitatively estimating filamentation difficult. From the mutants that could be quantitatively characterized at time zero, the *gcn4* mutant showed a statistically significant difference in filamentation (Holm corrected *t*-test, *p* < 0.05), but only in *C. dubliniensis* (Figure 2A). This mutant showed increased filamentation compared to the wildtype strain, although the difference was mild at this time point.

**Figure 2.**
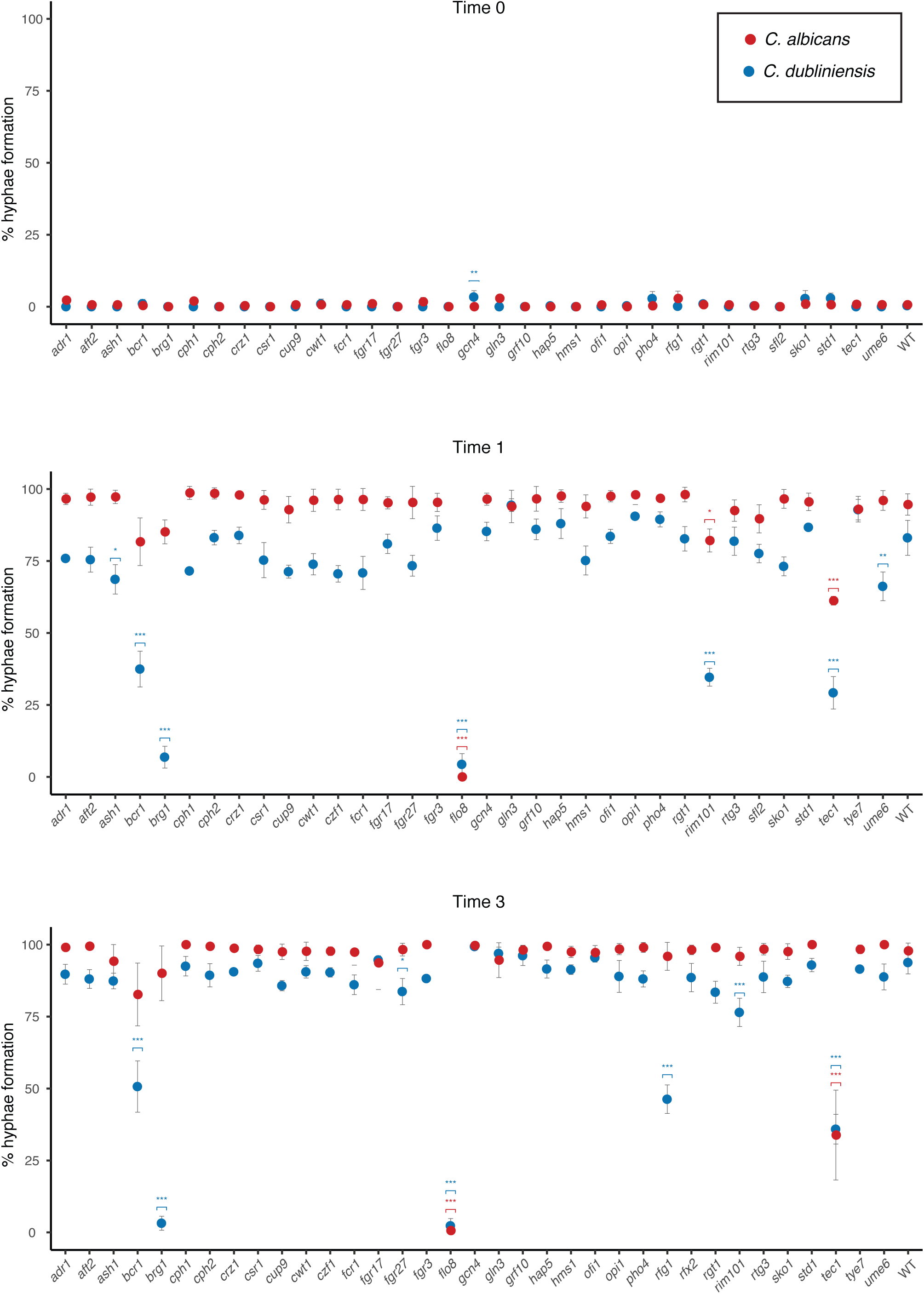
Multiple TR contribute to the filamentation differences between *C. albicans* and *C. dubliniensis*. Comparative phenotypic characterization of the TR mutants in filamentation inducing conditions (water with 10% FBS) through time. The percentage of the cells that showed germ tubes or hyphal morphologies is shown at each time point. Red dots represent the *C. albicans* mutants while blue ones the corresponding *C. dubliniensis* ortholog. Upper panel shows the phenotype when cells were transferred to the inducing conditions (Time 0) and the mid and lower panels show the phenotype after one (Time 1) and three (Time 3) hours after induction, respectively. Errors bars are the standard deviation of three replicates and only one of the isolates of each *C. dubliniensis* mutant is shown. Asterisks denote statistically significant differences between the mutant and the wild type strain (Holm corrected *t*-test, * *P* < 0.05, ** *P* < 0.01, *** *P* < 0.001). Only mutants that could be quantitatively assessed are included in each of the panels.

After one hour under the filamentation inducing conditions, apart from the mutants described above that could not be quantified at the zero-time point, the *rfg1* mutant also showed a morphology in *C. albicans* that impeded quantification. From the mutants in which filamentation could be quantified, *flo8, rim101* and *tec1* showed a statistically significant reduction in the number of filamenting cells in both species (Holm corrected *t*-test, *p* < 0.05). On the other hand, the mutants of *ASH1, BCR1, BRG1,* and *UME6* showed a statistically significant reduction in filamentation, but only in *C. dubliniensis*.

At the three-hour time-point in filamentation conditions, the *flo8* and *tec1* mutants continued showing a filamentation defect in both species, but in *rim101* the defect was only statistically significant for *C. dubliniensis*. The filamentation defect of the knockout strains of *BCR1* and *BRG1* in *C. dubliniensis* persisted at this time point, but *ash1* and *ume6* did not show statistically significant differences anymore. The only mutant that showed a filamentation defect at the three-hour time-point that had not shown the phenotype before was *fgr27*, and it did so only in *C. dubliniensis*. Contrary to the one-hour time-point, after three hours, filamentation could be quantitated in the *rfg1* and *rfx2* mutants and only the former showed a statistically significant difference compared to the wildtype strain in *C. dubliniensis*. Overall, among the mutants for which filamentation could be quantitated there were two main phenotypes: strains that showed more germ tubes than the wildtype strain at time zero, and mutants that had reduced number of filamentous cells after one and three hours in the inducing conditions.

In total, including the mutants for which filamentation was not quantitated, 22 of the TR mutants (50%) showed a filamentation phenotype different from that of the wildtype strain in one of the two species at least in one of the time points analyzed. If we only consider *C. albicans,* 11 mutants (25%) showed a filamentation defect. Of these mutants, ten also showed a defect in *C. dubliniensis,* while there were ten strains that only showed a phenotype in *C. dubliniensis.* It is important to point out that we performed the screen under specific inducing conditions and that alternative filamentation stimuli may be needed to expose the filamentation phenotypes of the *C. albicans* mutants that did not show a filamentation defect in our assay. However, in summary, our results suggest that several TRs contribute to the differences in the way filamentation is regulated in *C. albicans* and *C. dubliniensis*.

### The difference in filamentation between the *C. albicans* and *C. dubliniensis bcr1* mutant is condition dependent

One of the mutants that showed marked differences in filamentation between *C. albicans* and *C. dubliniensis* was *bcr1* (Figure 2). In *C. dubliniensis* this strain showed a considerable reduction in the number of filamentous cells after one hour in the inducing conditions, and after three and five hours it exhibited few pseudohyphal filaments. On the other hand, the mutant of the ortholog in *C. albicans* showed a similar fraction of germ tubes than the wildtype strain at the three time points. The phenotype in *C. albicans* is consistent with previous work showing that *BCR1* is not required for hyphal formation in planktonic growth conditions (27, 28).On the other hand, this gene is known to be required for biofilm formation in both species and in other *Candida* species that are phylogenetically further apart (20).

To investigate whether the observed differences in the *bcr1* mutants of the two species were specific to the growth conditions employed in the screen, we performed filamentation assays using Lee’s medium as a defined basal medium for induction. It has been reported that to induce filamentation in *C. dubliniensis* a combination of temperature and pH shift in nutrient depleted media is required, which can be achieved transferring cells to Lee’s medium at 37 °C. In addition, unlike fetal bovine serum, this medium does not include any animal-based components (18, 29). As can be observed in Figure 3, in contrast to the phenotype in fetal bovine serum, the *C. albicans bcr1* mutant showed decreased filamentation in Lee’s medium after one hour. After three hours in this medium, the defect of the *C. albicans* mutant was milder but still statistically significant. *C. dubliniensis* did not filament as efficiently in this medium but the defect of the *bcr1* mutant was evident along the time points (Figure 3).

**Figure 3.**
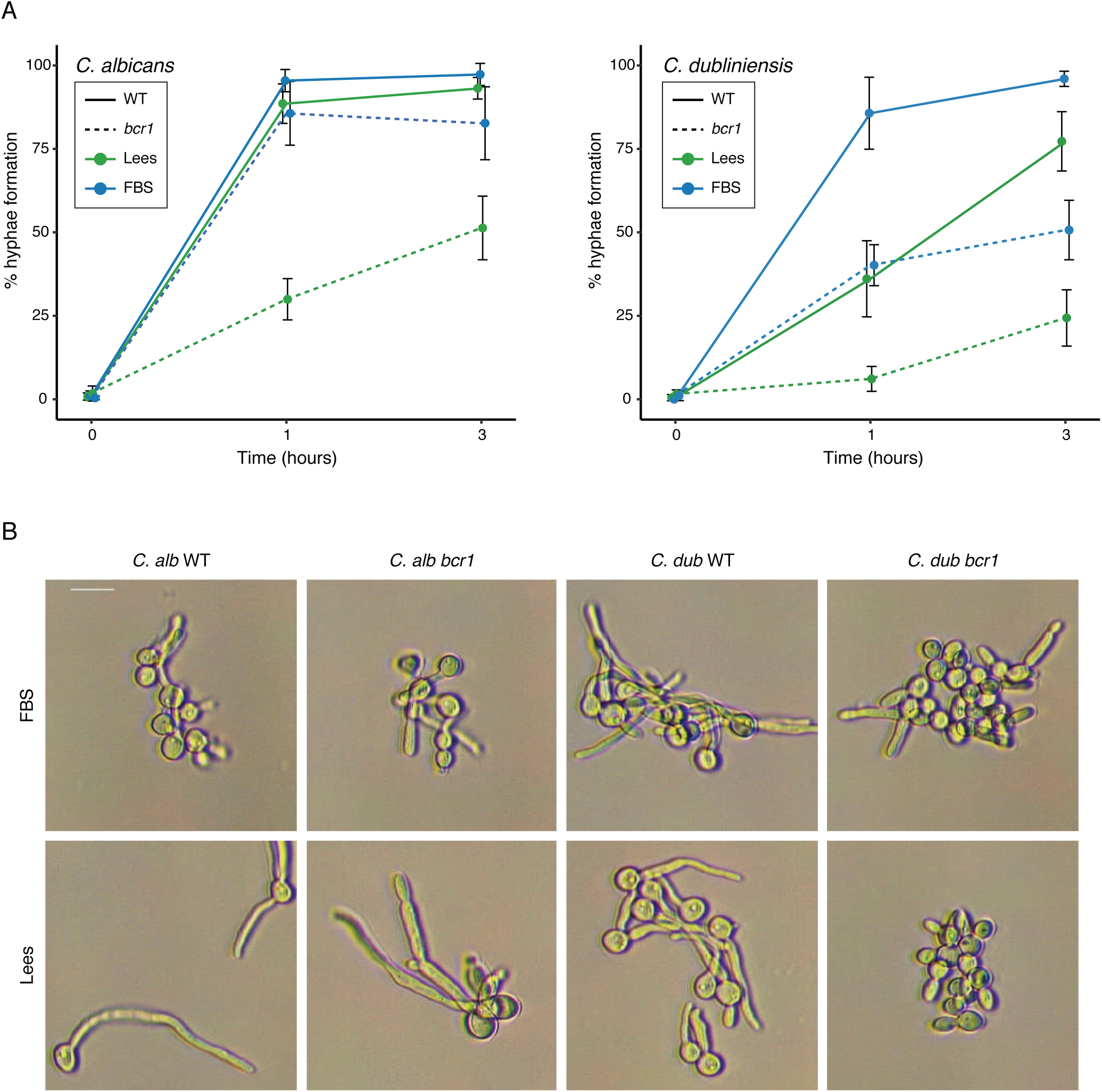
Filamentation differences of the *bcr1* mutant between species are media specific. (A) Quantification of the portion of cells that showed germ tubes or hyphal morphologies through time (0, 1 and 3 hours after induction) in the homozygous *bcr*1 mutant (discontinuous lines) compared with the wildtype strain (WT, continuous lines) in two different inducing media, water with 10% FBS (FBS, blue lines) and water with 10% Lee’s medium (Lees, green lines). Error bars show the standard deviation of three replicates. (B) Representative micrographs of the wildtype and *bcr1* mutant of the two species (*C. alb* and *C. dub*) filamenting in the same two media (FBS and Lees). Micrographs were taken under an optical microscope after three hours in the inducing condition and the scale bar represents 20 μm.

Given the results obtained when inducing by transferring to Lee’s medium, a condition previously reported to induce filamentation for *C. dubliniensis* (18), we examined the morphology of the *bcr1* mutants in response to other inducing cues, but that instead have been previously used for *C. albicans* (Materials and Methods). Of the three media tested, only SD supplemented with 0.75% glucose and 50% FBS induced filamentation in *C. dubliniensis*, and the *bcr1* mutant did not filament under this condition as was seen with FBS or Lee’s medium. The *C. albicans bcr1* mutant did not show a filamentation defect, behaving as the reference strain in the three additional media tested (Supplementary Figure 2).

We also assessed the role of the change in pH in the filamentation of the mutant, as it has been described as a crucial factor to induce filamentation in these species (18). For this, we set the pre-induction culture in Lee’s media adjusted at pH 7.2 instead of pH 4.5 as had been done for all previous assays. In this condition the *C. albicans bcr1* mutant showed an important filamentation defect not seen in the reference strain. This was observed when using FBS as the inducing condition, both in water or YPD medium. When starting at pH 7.5, the *C. dubliniensis bcr1* mutant showed the same defect as when the pre-induction culture was set at pH 4.5. Interestingly, when the pre-induction culture was set at pH 7.5 and RPMI medium supplemented with FBS was used for induction, the *C. dubliniensis bcr1* mutant did filament while the reference strain did not. Under this condition, both *C. albicans* strains filamented. Overall, these results showed that the filamentation requirement for Bcr1 depends on the conditions used to induce filamentation. However, it was also clear that *BCR1* is needed for filamentation in more conditions in *C. dubliniensis* than in *C. albicans*.

### Considerable differences in the gene expression program controlled by Bcr1 between ***C. albicans* and *C. dubliniensis***

To further understand the differences in the regulatory role of Bcr1 between *C. albicans* and *C. dubliniensis,* we focused on the genes that are controlled by this regulator during filamentation. We performed RNA-seq of the wildtype and the *bcr1* mutant in both species after one hour of filamentation induction with fetal bovine serum or Lee’s media. Comparison of the transcription profiles of the wildtype and *bcr1* mutant allowed us to identify the genes whose expression depends directly or indirectly on Bcr1 in the two different inducing conditions. Adding the two conditions and including all the genes whose expression change is statistically significant independently of the magnitude of the change, 2,198 and 2,296 genes were differentially expressed in the *bcr1* mutant in *C. albicans* and *C. dubliniensis*, respectively. This represents close to one third of the total number of genes in the genomes of these species. Comparing the expression profiles between filamentation conditions, we observed that the fraction of differentially expressed genes that are shared between media is larger in *C. dubliniensis* (39.8%) than in *C. albicans* (35.3%). This is in agreement with the observation that the filamentation defect was observed in both media in the mutant of *C. dubliniensis,* but not in that of *C. albicans*.

In the *bcr1* mutant, the overlap in differentially expressed genes between the two species was 16.5% in fetal bovine serum and 20.4% in Lee’s medium (Figure 4). This is also consistent with the fact that the filamentation phenotype of both species in Lee’s medium is similar, while it is contrasting in fetal bovine serum medium. The fraction of species-specific genes that are differentially expressed ranged from 6.3% in fetal bovine serum medium to 5.4% in Lee’s medium. These fractions are slightly smaller than the overall proportion of species-specific genes between these two species (9% of the total number of genes in the two genomes are species-specific (20)).

**Figure 4.**
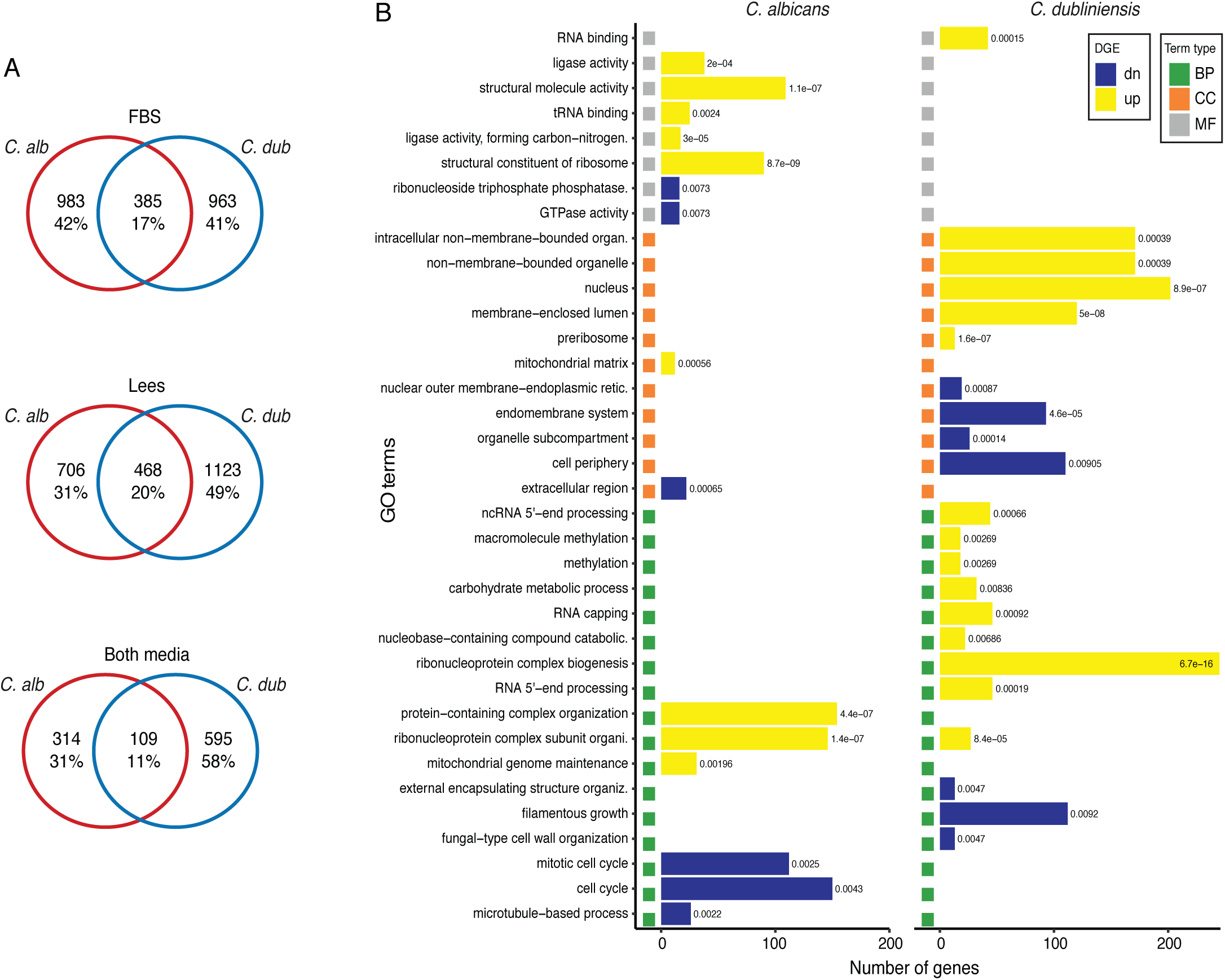
The transcription circuits controlled by Bcr1 have diverged considerably between *C. albicans* and *C. dubliniensis*. (A) Venn diagrams of the overlap between differentially expressed genes in the *bcr1* mutant of *C. albicans* and *C. dubliniensis*. The top diagram shows the overlap when inducing by transferring to water with 10% FBS (FBS), the middle diagram when inducing with water with 10% Lee’s medium (Lees), and the bottom diagram for differentially expressed genes in both media. All statistically significant differentially expressed genes were included, independently of the magnitude and direction of the expression change. (B) Comparison of the GO terms enriched among the differentially expressed genes in the two species. Only genes that were differentially expressed in both inducing conditions were included in the analysis. Yellow horizontal bars show categories enriched in upregulated genes while blue bars in downregulated genes. Colored squares at the left of each bar denote the type of GO term: BP, Biological Process; CC, Cellular Compartment; MF, Molecular Function. The numbers to the right of each horizontal bar are the enrichment *p*-value.

If we only consider the genes that are differentially expressed in the *bcr1* mutant in both inducing conditions, the fraction of gene targets that are shared between the two species is even smaller (10.7%, Figure 4). In total, *C. dubliniensis* had 727 differentially expressed genes across conditions, with 3.2% being species-specific. In *C. albicans*, the overlap between conditions was of 434 differentially expressed genes, of which 2.5% were species-specific. Performing GO enrichment analysis on the sets of genes that changed their expression in both conditions, we observed that the enriched categories are very different between the two species; only one category (*ribonucleoprotein complex subunit organization,* GO:0071826) out of 36 was enriched in both species (Figure 4). The category *filamentous growth* (GO:0030447) was only enriched in *C. dubliniensis* and for genes that are downregulated. This suggests that Bcr1 is needed to transcriptionally induce the filamentation program in this species. In *C. albicans* enriched functions in downregulated genes are instead related to the cell cycle (GO:0000278 and GO:0007049) and *microtubules-based process* (GO:0007017).

Focusing on the known TRs of filamentation in these species (Supplementary Table 3), we observed that the negative regulator Nrg1 is up regulated in both species and both conditions in the *bcr1* mutant. This is consistent with the notion that Bcr1 regulates this TR as it binds to its promoter region (30). The change in expression of *NRG1* is larger in *C. dubliniensis* when induced with fetal bovine serum (3.18 log2 vs 1.40 log2), but, conversely, in Lee’s medium *C. albicans* exhibits greater upregulation (1.66 log2 vs. 0.74 log2). These findings support the idea that in *C. dubliniensis* additional Nrg1-independent regulators are involved in repressing hypha formation when nutrients are present (16, 17). Interestingly, despite the overexpression of *NRG1* in the mutants of both species, the downstream filamentation regulator Ume6 was only downregulated in *C. dubliniensis* and in both inducing conditions. In *C. albicans* the expression of *UME6* is known to be repressed by Nrg1 and differences in its expression have been previously associated with the filamentation dissimilarities between the two species (17). Similarly, Brg1, another well-known regulator of filamentation in *C. albicans*, was downregulated in the *C. dubliniensis* mutant in both conditions and only slightly in serum for the *C. albicans* mutant. It has been suggested that hyphal development requires the upregulation of *BRG1* and the associated histone deacetylase Hda1, which together remodel the chromatin state of the hyphal-associated gene promoters and repress Nrg1 (31, 32). Tec1 was also downregulated in the *bcr1* mutant of *C. albicans* under both inducing conditions and more strongly in *C. dubliniensis* but not when inducing with Lee’s medium. This filamentation TR has been reported to be regulated by Bcr1, but also to feedback regulate *BCR1* expression (30).

Apart from *UME6* and in agreement with the enrichment of the GO category *filamentous growth* among the downregulated genes in *C. dubliniensis*, there were several genes previously associated with filamentation that were downregulated in this species in both conditions and that did not show an expression change in *C. albicans* (Supplementary Table 3). Among the best studied of these genes are *ALS1, DEF1* (both very strongly downregulated)*, CPH1* and *RIM101*. Similarly, *HWP1*, encoding a hyphal cell wall protein, was downregulated in the mutant of both species and under both conditions, but much stronger in *C. dubliniensis* (> 9 log2 fold in *C. dubliniensis* vs. < 1.7 log2 fold on *C. albicans*). In contrast, there were relatively few filamentation associated genes that were only upregulated in *C. dubliniensis*, such as *PHO4* and *RBF1*. The gene *RHD3* changed its expression considerably in the mutants of both species, but it did it in different directions.

There were also filamentation associated genes that only changed their expression in the *C. albicans* mutant in both inducing conditions, and most were downregulated (*RFX2, OPI1, ALS3, RBT5* and *ECE1*). The case of *ECE1* is interesting as this gene encodes Candidalysin, a hyphal-specific toxin that is needed for gut colonization (12). Among the most conspicuous genes due to the magnitude of their expression change even when they only changed in one of the inducing conditions were *SAP4* and *ALS2* in *C. albicans* and *SFL2* in *C. dubliniensis*, all three only changed in Lee’s medium. Overall, in addition to the specific expression differences between the two species in key filamentation regulators and genes such as Ume6, Brg1 and Tec1, our results revealed large scale contrasts in the transcription programs controlled by Bcr1 during filamentation in these two species.

## DISCUSSION

The morphological transition between yeast cells and hyphae is an essential trait for the colonization of the human body by pathogenic fungal species such as *C. albicans*. Dissimilarities in this transition could explain virulence differences between closely related species as has been suggested for *C. albicans* and *C. dubliniensis* (10). The wider range of environmental condition that have been observed to trigger filamentation in *C. albicans* seems to be associated with its higher clinical prevalence. At a molecular level, TRs such as Nrg1, Ume6 and Efg1 have been identified to underlie part of these differences (15, 17, 18). However, there are filamentation dissimilarities between these two species that seem to be independent of these TRs. To further understand the molecular underpinnings of the filamentation differences between *C. albicans* and *C. dubliniensis* we focused on all the TRs that had been previously implicated in filamentation in *C. albicans*, the most studied species. The orthologs of the 45 TRs in the two species are highly similar at the sequence level, containing the same protein domains. However, analysis of a few available DNA binding motifs that these TRs bind revealed extensive differences in their genomic distribution between species. These observations suggest that for these TRs and at the distance between *C. albicans* and *C. dubliniensis* (∼ 20 million years) most evolutionary changes occurred in the target genes that they control through *cis*-regulatory mutations. This agrees with previous observations in the transcription circuits that regulate biofilm formation and glycolysis in these two species (20, 33).

The general high sequence similarity between ortholog TRs and low conservation of the distribution in DNA binding motifs also showed that without experimentation it would be difficult to pinpoint further TR that are responsible for the filamentation differences between *C. albicans* and *C. dubliniensis*. For this reason, we generated a collection of knockout mutants of the *C. dubliniensis* orthologs of the 45 *C. albicans* filamentation TRs. The only TR for which we could not delete both alleles was Tup1, despite several knockout attempts. This was surprising since there are reported mutants of this TR in *C. albicans* and *C. tropicalis*, the two most closely related species (22, 23). In *C. albicans,* Tup1 is a known repressor of filamentation genes by interacting with the DNA binding TR Nrg1 (4). One of the explanations for our inability to knockout *TUP1* in *C. dubliniensis* is that this gene has gained a more general role in this species and has thus become essential. Further work will be needed to elucidate the role of *TUP1* in *C. dubliniensis* filamentation and its overall cellular physiology.

Functional characterization of the *C. dubliniensis* gene-knockouts in parallel with the corresponding *C. albicans* mutants revealed several TRs with contrasting filamentation phenotypes, beyond previously known differences (Figure 2 and Supplementary Figure 1). Validating our approach, we observed phenotypic interspecific differences in the *nrg1* and *ume6* mutants, the two TRs that have been previously more strongly associated with the filamentation dissimilarities between these species (15, 17). On the contrary, we did not observe differences in the *efg1* mutants. This TRs has been proposed to regulate several filamentation genes that are only differentially expressed in *C. albicans* (18). However, the role of this TR in biofilm formation is known to be conserved between the two species (20), which would agree with the conservation that we observed in terms of the filamentation defects of the *efg1* mutants in both species (Supplementary Figure 1).

Among the most studied TRs that, to our knowledge, had not been previously associated with filamentation differences between the two species are *BCR1* and *BRG1*; the mutants of these TRs showed reduced filamentation only in *C. dubliniensis* under the inducing conditions of the screen. In agreement, Bcr1 is known not to be required for filamentation in white *C. albicans* cells, but to be a repressor of this morphological transition in opaque cells (30). In contrast, Brg1 is a known regulator of filamentation in the white *C. albicans* state, while it does not seem to be important for hype formation in opaque cells (30). Furthermore, both of these TRs are central for biofilm formation in both species (20). Other mutants with marked differences included *rca1* whose phenotype is similar to that of the reference strain in *C. dubliniensis*, while it was hyperfilamentous in *C. albicans*, resembling the *nrg1* mutant. Deletion of *STP2* was also contrasting as the mutant did not filament in *C. dubliniensis* while it did in *C. albicans,* although forming unusually thick filaments and cells. Overall, our screen showed that several TRs are responsible for the differences in filamentation between these two species and suggest key TRs for future work.

Bcr1 was originally described as a zinc finger TR needed for biofilm formation in *C. albicans*, but that was dispensable for filamentation (28). This was surprising given that biofilm formation and filamentation are tightly interconnected processes. *BCR1* was latter associated with filamentation, although only in opaque cells (30). Therefore, we did not expect to observe a filamentation defect in the *bcr1* mutant of *C. dubliniensis,* as we did. Further characterization of the *bcr1* mutants in a variety of inducing media showed that the filamentation defect was condition specific. We found media in which the *C. albicans* mutant did show a filamentation defect, but also in which the *C. dubliniensis* knockout was able to filament (Figure 3, Supplementary Figure 2). The general observed trend, however, is that there were more inducing conditions in which *C. albicans* mutant did not show a phenotype when compared to the *C. dubliniensis* mutant.

Transcriptional profiling in two different inducing conditions showed considerable interspecific differences in the genes that changed their expression in the *bcr1* mutant (Figure 4). This holds true even if we only take into account the genes that showed a larger expression change. In fact, using a 1.5 or 2 log2 fold change cutoff reduced the commonly differentially expressed genes between the two species. There are not many studies were genome-wide gene expression has been compared between *C. albicans* and *C. dubliniensis,* and most compare profiles done in different laboratories (17, 18). However, the overall degree of conservation from experiments done in a single study is not far from what we observed. For example, between 22 and 29% of differentially expressed genes during biofilm formation in these two species was reported to be conserved (20), while we observed 17 and 20% in the two inducing conditions here tested. These results are also consistent with the low degree of conservation in the genomic distribution of the binding motifs of the analyzed TR. At this point it is difficult to say whether the overlap between the transcription programs controlled by Bcr1 in the two species is lower than expected and could underly the interspecific filamentation differences observed in the mutants.

The transcription profiles reveled several filamentation genes that are controlled by Bcr1 and that could explain the interspecific filamentation differences in the mutants of this TR. Interestingly, *UME6* seems to be activated by Bcr1 only in *C. dubliniensis*. This gene encodes a TR that is an important activator of the filamentation program and differences in its expression have been previously proposed to partially underly the hypha formation dissimilarities between *C. albicans* and *C. dubliniensis* (9, 16). *UME6* is known to be negatively regulated by the filamentation repressor Nrg1 and considerable emphasis has also been placed in this other regulator to explain filamentation in these species. Our experiments revealed that *NRG1* is repressed by Bcr1 in both species, although more strongly in *C. dubliniensis* when induced with FBS. Several other filamentation genes had a similar expression pattern than *UME6,* being down regulated only in the *C. dubliniensis bcr1* mutant. In fact, the functional category *filamentous growth* was only enriched in downregulated genes in the mutant of *C. dubliniensis*, suggesting that in this species Bcr1 plays a more central role regulating filamentation. Still, in *C. albicans*, Bcr1 specifically performed as a positive regulator of key hyphal factors such as *ECE1*.

In summary, our work showed that several TRs, beyond previously known ones, seem responsible for the filamentation differences between *C. albicans* and *C. dubliniensis*. In addition, the role of these transcription circuits is condition dependent reflecting the complexity of the filamentation programs in these species. We propose that Bcr1 plays a more predominant role in the filamentation of *C. dubliniensis*, controlling the expression of key TRs such as Ume6. Overall, the degree of dissimilarities found between *C. albicans* and *C. dubliniensis* at a molecular level may not be that surprising if we consider that we are comparing organisms that diverged at least as early as humans and gibbons did. The results of our filamentation screen and the collection of gene-deletion mutants will be valuable resources to further understand the virulence differences between *C. albicans* and *C. dubliniensis* in the near future.

## MATERIALS AND METHODS

### Identification of TRs associated with filamentation

TRs associated with filamentation in *C. albicans* were identified from the information available at the Candida Genome Database (CGD) (34). A TR, as defined by Homann O. R., *et al*. (2009) (21), was considered associated with filamentation if it was part of the Phenotype Term “filamentous growth” including the following phenotypes: “filamentous growth: abnormal”, “filamentous growth: increased”, “filamentous growth: decreased”, “filamentous growth: absent”, “filamentous growth: decreased rate” and “filamentous growth: delayed”. This is a similar strategy to what has been used by Noble, S. M., *et al*. (2010) (35) to identify virulence associated genes. *C. dubliniensis* one-to-one orthologs were identified from the orthology assignments at the Candida Gene Order Browser (19).

### Estimation of the amino acid sequence identity between TR orthologs and identification of protein domains

*C. albicans* (C_albicans_SC5314_version_A22-s07-m01-r177_default_protein) and *C. dubliniensis* (C_dubliniensis_CD36_version_s01-m02-r36_orf_trans_all) TRs amino acid sequences were obtained from CGD to be then pairwise aligned using MUSCLE with default parameters (36). DNA binding domains of each TR were identified with the InterProScan 5.57-90.0 motif finding algorithm (37) and the sequence of the motifs of both species was aligned also using MUSCLE.

### Computational determination of putative target genes

Empirically determined DNA binding motifs of *C. albicans* TRs were obtained from PathoYeastract (Pathogenic Yeast Search for Transcriptional Regulators And Consensus Tracking) (38). These motifs were used to scan the *C. albicans* (C_albicans_SC5314_version_A22-s07-m01-r109_chromosomes) and *C. dubliniensis* (C_dubliniensis_CD36_version_s01-m02-r26_chromosomes) intergenic regions using MochiView (39) with standard parameters. Putative target genes were defined as those having at least one motif in the 2 kb upstream intergenic region. If there was another ORF closer than 2 kb, the limit of the neighboring ORF was set as the end of the intergenic region.

### Generation of the knockout mutant collection of filamentation TRs in *C. dubliniensis*

Homozygous null mutants of the *C. dubliniensis* TRs were generated following the genetic modification strategy previously described by Mancera E., *et al.* (2019) (40). In brief, fusion PCR was performed to generate *HIS1* and *LEU2* gene disruption cassettes. To this end, *HIS1* and *LEU2* nutritional markers were amplified from the plasmids pEM001 and pEM002, respectively, with primers 2 and 5. In parallel, approximately 350 nucleotides of the flanking downstream and upstream sequences of the ORF to be deleted were amplified from genomic DNA of *C. dubliniensis* strain CD36 with primers 1 - 3 and 4 - 6 in separated reactions. The nutritional markers were then stitched together with the up and downstream homology regions by fusion PCR with primers 1 and 6. To delete the first allele and generate the heterozygous mutant, the *HIS1* disruption cassette was transformed in parallel into CEM074 and CEM075 auxotrophic strains by electroporation (40). The homozygous deletion strain of each TR gene was then generated by transformation of the heterozygous strain with the *LEU2* disruption cassette. Selection of the transformants was performed by growing cells on synthetic defined (SD) medium (6.7 g/L yeast nitrogen base, 2% glucose, supplemented with amino acids) without histidine or leucine. Verification of the correct integration of the deletion cassettes was performed by colony PCR of the 5’and 3’ junctions. In addition, after deleting the second allele, the absence of the target gene was verified with primers that amplify a region within the targeted ORF. Two knockout strains were constructed for each *C. dubliniensis* gene, one in each of the two parental strains CEM074 and CEM075 (40). Strains used in this study are listened in Supplementary Table 2.

### Screening the *C. albicans* and *C. dubliniensis* mutant collections for filamentation defects

*C. albicans* and *C. dubliniensis* strains were streaked out from glycerol stocks in SD medium without histidine and leucine. A single colony from these plates was used to start liquid cultures in Leés medium adjusted to pH 4.5 (17, 29). These cultures were grown at 30 °C for 18 hrs. The cells were then washed twice with Milli Q water. Filamentation was induced by inoculating 2 x 10^6^ cells to Milli-Q H2O supplemented with 10% (v/v) fetal bovine serum (17, 18) on 24-well cell culture plates. Cultures were incubated at 37 °C shaking at 200 rpm. The percentage of germ tubes or filamentous cells was counted under an optical microscope at the time of transferring to inducing condition, and one and three hours after. Three replicate assays were performed for each of the two *C. dubliniensis* mutants of each TR. A Holm multiple corrected *t*-test was used to compare the percent of hypha formation in each TR mutant with that of the reference strain in each species at each time point. At least ten different view fields were counted per strain, containing a minimum of eight cells per field. For each filamentation TR two independent mutant strains were characterized for *C. dubliniensis* and one for *C. albicans*. Strains CEM091 and CEM092 were used as a wildtype reference for *C. dubliniensis* (40), and SN250 for *C. albicans* (35).

To characterize filamentation of the *bcr1* mutants of both species in a wider set of conditions, cultures were grown in Lee’s medium at pH 4.5 or pH 7.2 incubating at 30 °C for 18 hours with constant agitation (17, 18). Filamentation was then induced by inoculating 2 x 10^6^ cells in 24-well cell culture plates and incubating at 37 °C with shaking at 200 rpm. The following nutrient-rich media were used to evaluate filamentation capacity: YPD and RPMI 1640, both supplemented with 10% fetal bovine serum (FBS, v/v) (8, 17), and SD medium with 0.75% glucose, supplemented with 50% FBS. The latter medium has been reported to be optimal to induce filamentation in *C. tropicalis* (41). Low-nutrient filamentation inducing media employed were Milli-Q water supplemented with 10% FBS or Lee’s medium adjusted to pH 7.2 (18).

### Transcriptional profiling

For RNA extraction, strains were grown in Leés medium pH 4.5 at 30 °C for 18 h. Cells were then washed twice in Milli Q water and approximately 2 x 10^6^ cells were inoculated in the inducing medium and incubated at 37 °C for one hour shaking at 200 rpm (18). Inducing media employed was Milli-Q H2O supplemented with 10% (v/v) fetal bovine serum or 10% (v/v) Leés glucose medium pH 7.2 as for the filamentation assays described above. Total RNA was extracted using the RiboPure™-Yeast Kit (Ambion) with some minor modifications. Cells were collected by centrifugating cultures at 3500 rpm for 5 minutes, the lysis components (Lysis Buffer, 10% SDS and Phenol:Chloroform:IAA) were immediately added to the pellet and they were then frozen at -80 °C. The frozen pellets were thawed on ice and the rest of the protocol was performed as indicated by the manufacturer but performing all steps at 4 °C in a cold room. Three replicates were performed for each strain (wildtype and *bcr1* mutant) in each inducing medium for each species. Quality control of the total RNA, library preparation and sequencing were performed by Novogene Co. All samples had a RIN higher than 4.0. Libraries were directional using poly A enrichment and sequencing was PE150 in a NovaSeq platform. A minimum of 2 G raw data per sample were obtained.

To identify differentially expressed genes, raw reads were first cleaned with fastp v0.46.2 (42) to remove read regions of low quality, potential adaptor sequences, poly(A)-tails and long terminal homopolymeric stretches. Clean reads were then aligned and quantified using kallisto v0.46.2 (43) against the cDNA transcripts reported in CGD. Strains SC5214 (A22-s07-m01-r168_default_coding) and CD36 (s01-m02-r36_orf_coding) were used as reference for *C. albicans* and *C. dubliniensis*, respectively. To identify anomalous samples, an outlier map was made based on the robust score distances and orthogonal distances computed by the PCAGrid function (44) for the normalized count matrix of each dataset as suggested in (45). Based on these results, two replicates of *C. dubliniensis* in Lee’s medium (one for the wild-type strain and one for the *bcr1* mutant) and three replicates of *C. albicans* (one of the *bcr1* mutant in FBS medium, one of the wild-type strain in Lee’s medium, and one of the *bcr1* mutant in Lee’s medium) were removed. A cutoff of 97.5% of the corresponding distribution was used to classify the samples. Finally, the exactTest function of the edgeR package (46) was used to determine gene differential expression. The resulting *p*-values were corrected with the *q*-value function using the default parameters to obtain an FDR of 1%.

The list of differentially expressed genes for each species was then used as input for a GO enrichment analysis using the R package topGO (47). GO terms were mapped to genes using the readMappings function and gene-to-GOs association data provided by CGD. Enrichment analysis was performed with Fisher’s exact test, using the classic algorithm, and a significance cutoff of *p*-value < 0.01. To summarize the enriched GO terms, we utilized the REVIGO online analysis tool (48), allowing a similarity score of 0.7 between GO terms. The UniProt GO database was used for mapping, and SimRel was selected as the semantic similarity measure.

## DATA AVAILABILITY

The raw RNAseq data have been deposited at the NCBI under BioProject ID PRJNA1163420.

## ACKNOWLEDGEMENTS

We thank Susana Ruiz-Castro, Fernando Villanueva Rodríguez, Silvia Lisset Juarez Valtierra, and Claudia Geraldin León-Ramírez for technical assistance.

## AUTHOR CONTRIBUTIONS

Conceptualization: EM. Methodology: TM-D, LFG-O. Investigation, laboratory work: TM-D. Investigation, formal analysis: TM-D, LFG-O, EM. Visualization: TM-D, LFG-O, EM. Funding acquisition: EM. Supervision: EM. Writing, original draft: TM-D, EM. Writing, review and editing: LFG-O. All authors read and approved the final version of the manuscript.

## MANUSCRIPT VERSIONS

This manuscript was released as a pre-print at *bioRxiv* (Meza-Davalos T., et al., 2024).

---

**Supplementary Figure 1.**
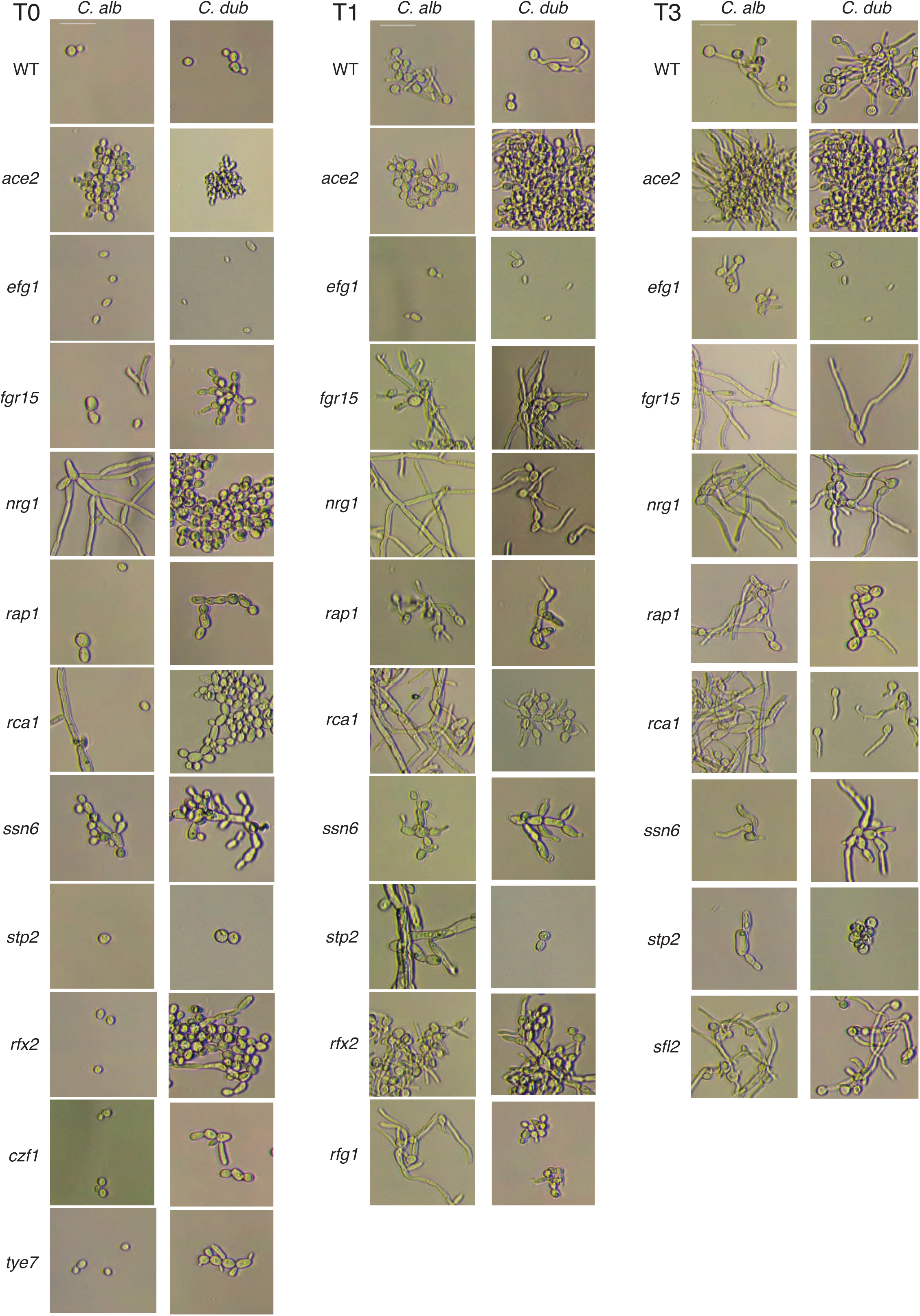
Phenotypes of the mutants for which filamentation could not be quantified. Micrographs taken under an optical microscope at the time the cells were transferred to the filamentation inducing conditions (T0) and after one (T1) and three hours (T3) of filamentation. The *C. albicans* (*C.alb*) and *C. dubliniensis* (*C. dub*) homozygous mutants are shown side by side. Only one of the *C. dubliniensis* isolates is shown although the phenotype was similar in the other isolate. The reference scale bar in the wildtype *C. albicans* represents 10 μm.

**Supplementary Figure 2.**
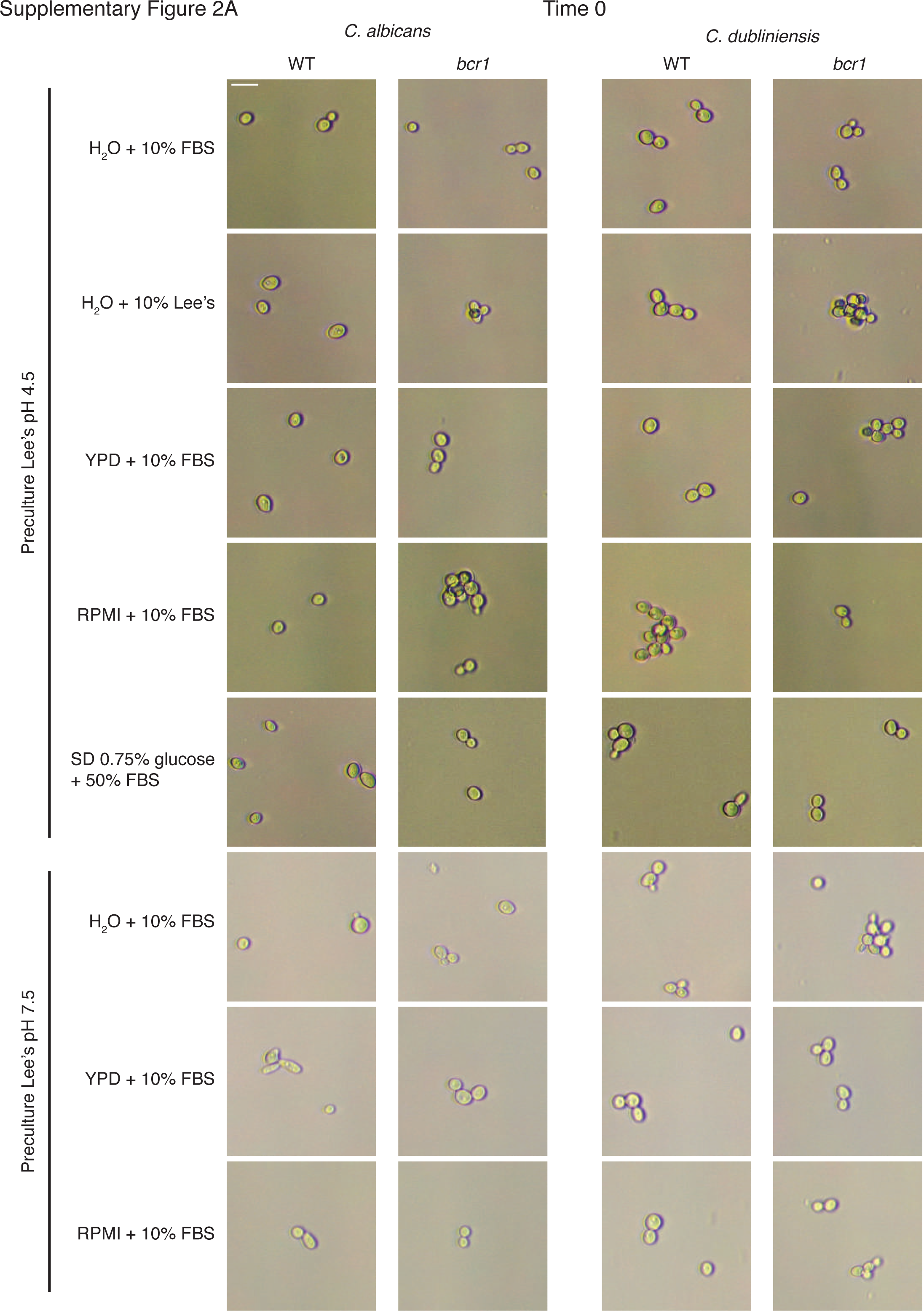
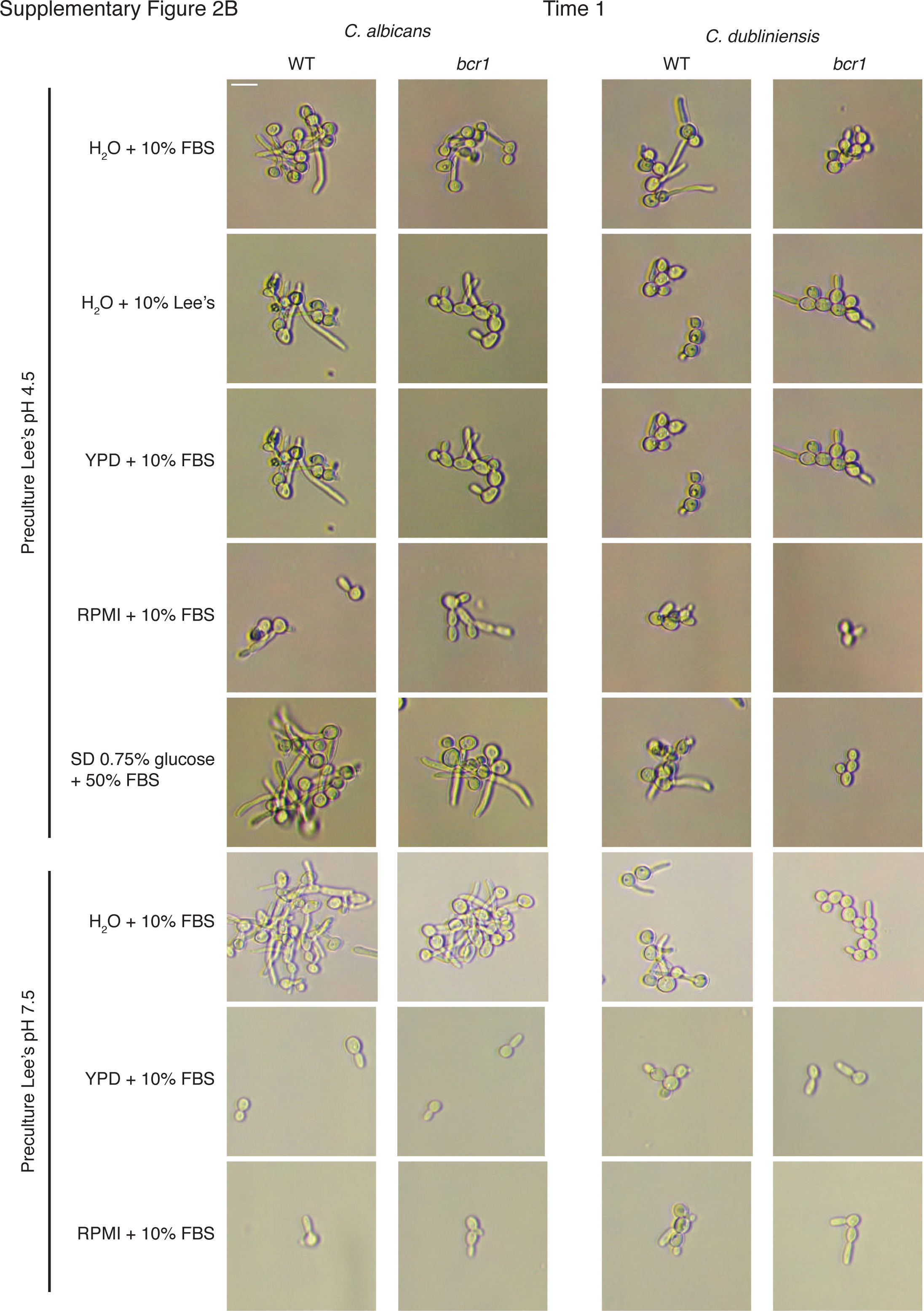
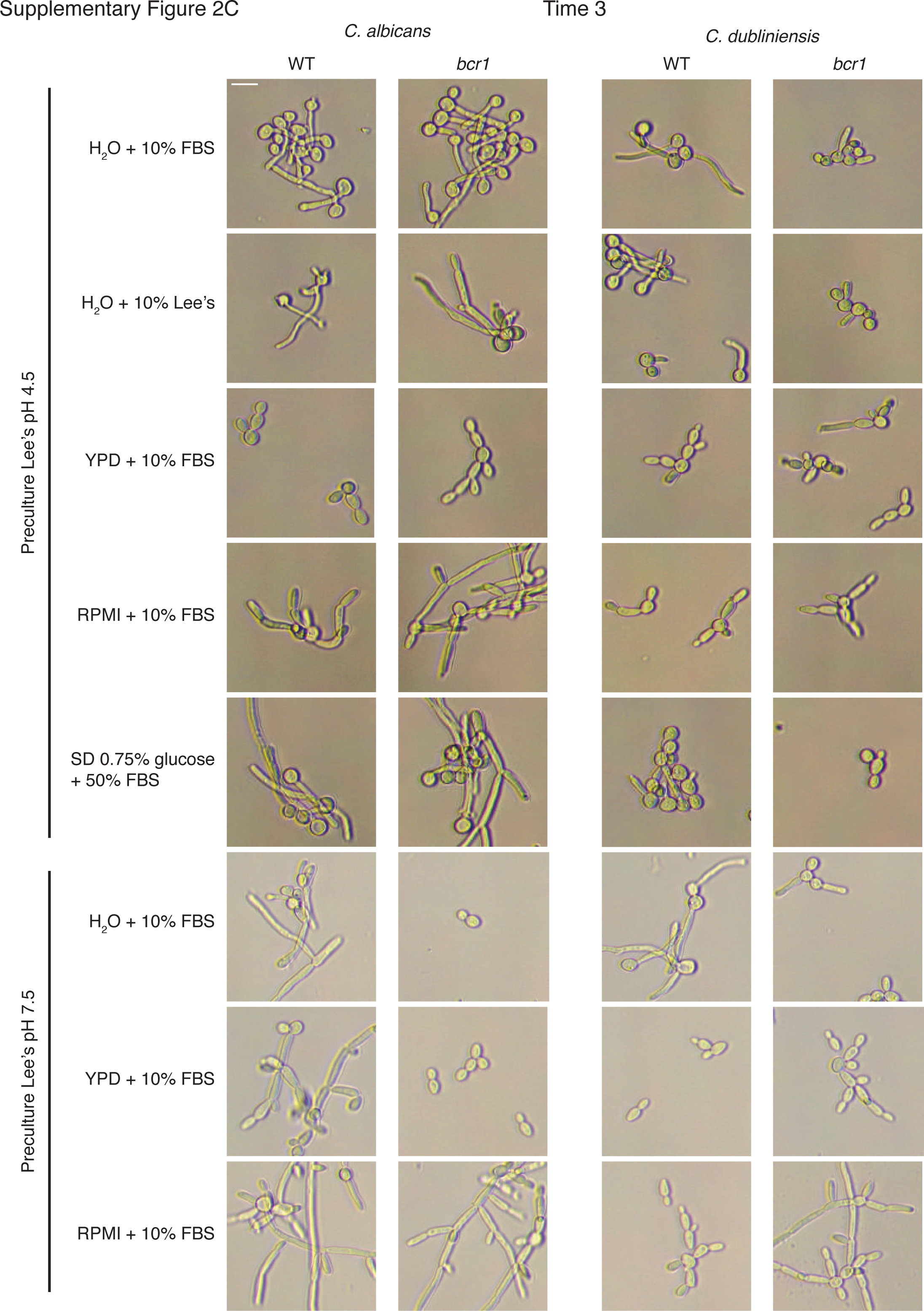
Morphology of the *bcr1* mutant under different filamentation inducing conditions. Micrographs under a light microscope of the reference strain and the *bcr1* mutant of *C. albicans* and *C. dubliniensis* in the different media tested. As indicated at the furthest left, two preculture conditions were employed. Panel A, B and C show micrographs taken at the 0, 1 and 3 hour time points, respectively. The reference scale bar in the wildtype *C. albicans* in H20 + 10% FBS when preculture in Lee’s pH4.5 was used represents 10 μm.

## SUPPLEMENTARY TABLES

**Supplementary Table 1.** *C. albicans* filamentation TRs considered in this study.

**Supplementary Table 2**. Yeast strains used and generated in this study.

**Supplementary Table 3**. Expression of genes related to filamentation in the *bcr1* mutants of *C. albicans* and *C. dubliniensis*.

